# Proteome-wide reverse molecular docking reveals folate receptor as a mediator of PFAS-induced neurodevelopmental toxicity

**DOI:** 10.1101/2024.11.11.623082

**Authors:** Ally Xinyi Kong, Maja Johnson, Aiden F Eno, Khoa Pham, Ping Zhang, Yijie Geng

## Abstract

Per- and polyfluoroalkyl substances (PFAS) are a class of long-lasting chemicals with widespread use and environmental persistence that have been increasingly studied for their detrimental impacts to human and animal health. Several major PFAS species are linked to neurodevelopmental toxicity. For example, epidemiological studies have associated prenatal exposure to perfluorooctanoate (PFOA) and perfluorononanoate (PFNA) with autism risk. However, the neurodevelopmental toxicities of major PFAS species have not been systematically evaluated in an animal model, and the molecular mechanisms underlying these toxicities have remained elusive. Using a high-throughput zebrafish social behavioral model, we screened six major PFAS species currently under regulation by the Environmental Protection Agency (EPA), including PFOA, PFNA, perfluorooctane sulfonate (PFOS), perfluorohexanesulfonic acid (PFHxS), perfluorobutane sulfonate (PFBS), and hexafluoropropylene oxide dimer acid ammonium salt (GenX). We found that embryonic exposure to PFNA, PFOA, and PFOS induced social deficits in zebrafish, recapitulating one of the hallmark behavioral deficits in autistic individuals. To uncover protein targets of the six EPA-regulated PFAS, we screened a virtual library containing predicted binding pockets of over 80% of the 3D human proteome through reverse molecular docking. The screen predicts that folate receptor beta (FR-β, encoded by the gene *FOLR2*) interacts strongly with PFNA, PFOA, and PFOS but to a lesser degree with PFHxS, PFBS, and GenX, correlating positively with their *in vivo* toxicity. These predictions were validated through *in silico* molecular docking, *in vitro* protein binding analysis, and *in vivo* loss-of-function verification. Furthermore, embryonic co-exposure to folic acid effectively rescued social deficits induced by PFAS. The folate pathway has been implicated in autism, indicating a novel molecular mechanism for PFAS in autism etiology.

## INTRODUCTION

Autism spectrum disorder (ASD) is a complex neurodevelopmental condition marked by social communication deficits. Over recent decades, the prevalence of autism has risen significantly, with reported cases in the United States increasing from 1 in 150 children diagnosed in 2000 to 1 in 36 in 2020, according to the Centers for Disease Control and Prevention (CDC). This increase is only partially accounted for by improved diagnostic criteria and awareness^1,2^. Meanwhile, the ever-growing chemical industry continues to introduce new chemicals into our environment, most of which with unknown toxicity to humans. An estimated 10 million new chemicals are synthesized annually, and within them, ∼ 700 become mass produced for commercial use^3^. Emerging research indicates that environmental factors contribute substantially to autism risk, with recent estimates attributing up to 40% of ASD etiology to environmental influences^4,5^. Understanding how environmental factors contribute to ASD is critical to mitigating the disease prevalence and could inform targeted prevention and treatment strategies.

However, environmental contributors to ASD remain poorly understood, creating a gap in research methods and assessment approaches to identify and evaluate these risk factors comprehensively. Among environmental toxins with known associations with ASD, per- and polyfluoroalkyl substances (PFAS) have become a particular focus of recent research^6,7^. PFAS is a class of synthetic chemicals known for their persistence in the environment and biological systems. They are widely present in nonstick cookware, stain-resistant textiles, and various industrial applications. Their pervasive use has led to widespread human exposure, with the CDC reporting PFAS detected in the blood of 97% of Americans. The health risks associated with PFAS exposure are substantial, including links to kidney and liver disease, immune and thyroid dysfunction, cancer, and, critically, developmental challenges^8^. Based on these pioneering studies, beginning in April 2024, the U.S. Environmental Protection Agency (EPA) established legally enforceable levels for six PFAS in drinking water, including perfluorooctanoic acid (PFOA), perfluorooctane sulfonic acid (PFOS), perfluorononanoic acid (PFNA), perfluorohexanesulfonic acid (PFHxS), perfluorobutane sulfonate (PFBS), and hexafluoropropylene oxide dimer acid ammonium salt (GenX). Despite these progresses, however, with 15,538 PFAS compounds identified in the Distributed Structure-Searchable Toxicity (DSSTox) database by the EPA^9^ and over 7 million PFAS species reported in PubChem to date^10^, a comprehensive understanding of their biological impact remains elusive, particularly regarding how prenatal exposures to various PFAS species contribute to ASD risk. For example, although epidemiology studies linked PFNA and PFOA to increasing ASD risk^6,7,11^, whether the other four EPA-regulated PFAS contribute equally to ASD remains unclear. Mechanisms underlying their neurodevelopmental toxicity are also largely unknown.

To address these gaps, we developed a novel model and methodology using zebrafish. Zebrafish offer a unique advantage as a model organism for studying social behavior^12^—a core behavioral deficit in ASD. They share approximately 70% of their genome with humans, begin social interactions at an early developmental stage, and are amenable to high-throughput chemical screening and behavioral assays. In our previous work, we designed the Fishbook social behavioral assay^13^ to quantitatively assess social behavioral deficits in zebrafish, a behavior analogous to social deficits in ASD. In this assay, a test fish is exposed to a social stimulus in a controlled environment, allowing us to measure a social score based on proximity behaviors. Notably, the Fishbook assay has observed social deficits in zebrafish following ASD-related gene knockouts and exposure to ASD-related environmental toxicants^13–15^, indicating its relevance for examining environmental toxicants’ influence on ASD. The Fishbook system enables us to screen different PFAS compounds in parallel using the same assay platform for their impacts on sociality, alongside genetic knockout models to elucidate their molecular mechanisms.

Moreover, to uncover potential molecular targets of PFAS, we conducted virtual screening through reverse molecular docking using the HPRoteome-BSite database^16^ to explore interactions between PFAS and the predictable 3D human proteome. HProteome-BSite is a database of predicted binding pockets covering over 80% of the UniProt entries in the AlphaFold human proteome database^17,18^. The reverse molecular docking approach, unlike traditional docking methods which typically screen a large number of chemicals against one or a few specific proteins, allows us to comprehensively evaluate how different PFAS species interact with a range of proteins involved in neurodevelopmental pathways.

In this study, we leverage our unique zebrafish model and assay with advanced virtual screening techniques and validation experiments to evaluate the impacts of the six EPA-regulated PFAS on social behavioral development and identified their molecular targets in the human proteome. Among the molecular targets predicted by the virtual screen, folate receptors were found to differentially bind to the six PFAS species, with the binding affinities positively correlating with their toxicity on social development. *In silico*, *in vitro*, and *in vivo* experiments verified folate receptor beta as a biological target of PFNA, PFOA, and PFOS. Co-exposure to folic acid rescued social deficits caused by PFAS. Given the importance of folic acid in early neurodevelopment^19–21^ and its implications for autism^22^, our findings strongly suggest that PFAS’ interaction with folate receptors likely contribute to their ASD-related neurodevelopmental toxicity and point to a potential strategy for preventive interventions.

## RESULTS

### The six EPA-regulated PFAS showed differential impacts on social behavioral development in zebrafish

To compare the impacts of major PFAS species on social behavioral development, we tested the six EPA-regulated PFAS, namely PFNA, PFOA, PFOS, PFHxS, PFBS, and GenX (Figure 1A), using the Fishbook assay (Figure 1B). After an initial embryonic toxicity testing, a range of doses were determined for each PFAS to assess their dose responses. The highest doses for the dose-response analysis were set as 50% of the lowest doses that caused 100% mortality in the initial toxicity test. The remaining doses for the dose-response analysis were set as 50% serial reductions from the highest dose. Specifically, PFNA was evaluated at 0.78 µM to 200 µM, PFOA at 3.125 µM to 1600 µM, PFOS at 0.25 µM to 12 µM, PFHxS at 3.125 µM to 400 µM, PFBS at 3.125 µM to 1600 µM, and GenX at 1.56 µM to 1600 µM. Embryos were exposed without dechorionation, with no observed impacts on hatchability due to PFAS exposures. Embryos were exposed to a single dose of PFAS at the concentrations listed above from 3 hours post fertilization (hpf) to 3 days post fertilization (3 dpf).

**Figure 1.**
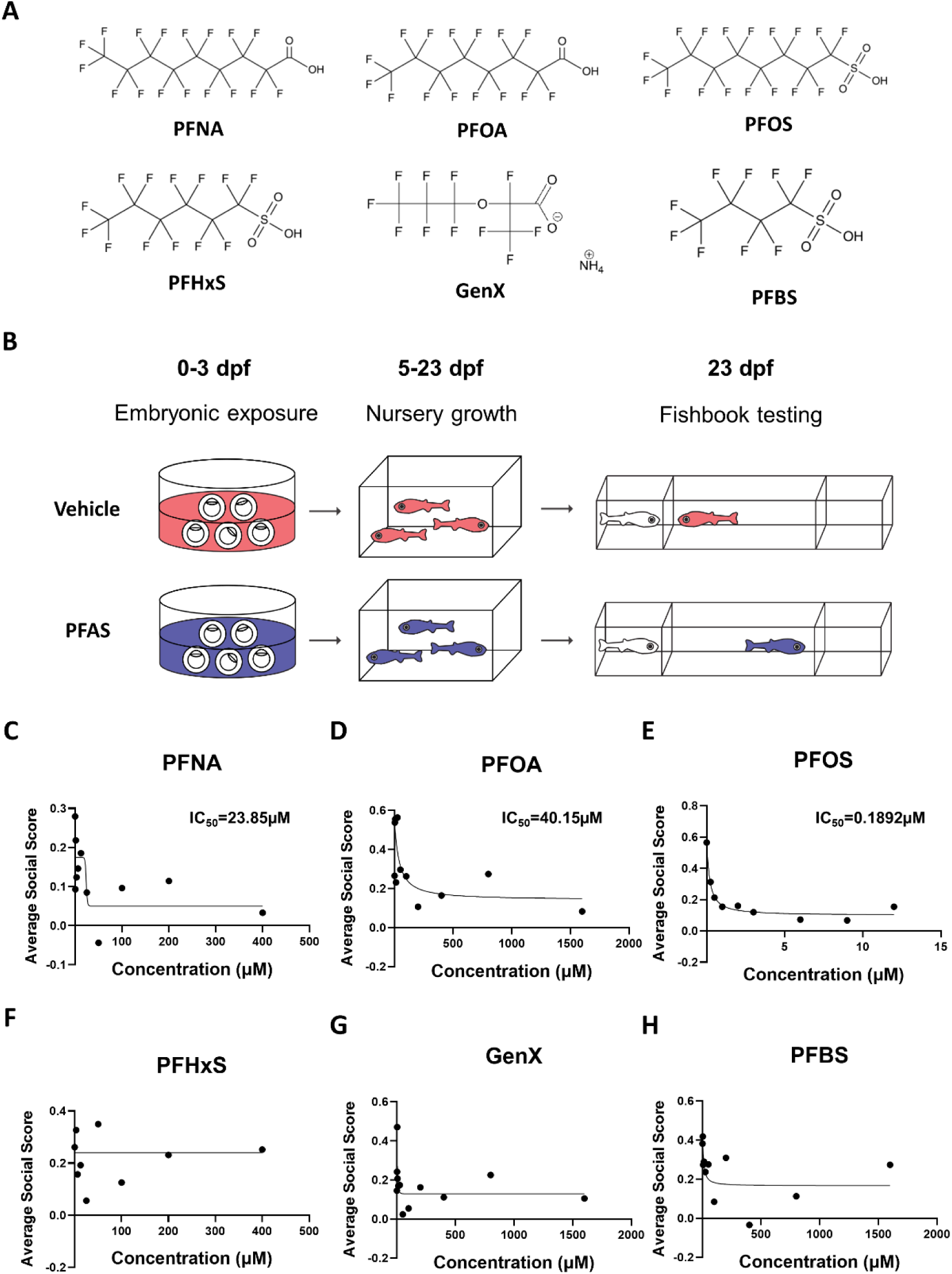
Fishbook assay revealed differential neurodevelopmental toxicity of six EPA-regulated PFAS on social behavioral development in zebrafish. **(A)** Chemical structures of the six EPA-regulated PFAS. (**B**) A schematic of the dosing and testing workflow. (**C-H**) Dosage curves generated by plotting average social scores (y-axis) against PFAS concentrations (x-axis). PFAS tested include PFNA (C), PFOA (D), PFOS (E), PFHxS (F), GenX (G), and PFBS (H). IC50 values are shown for PFNA (C), PFOA (D), and PFOS (E). Sample sizes are as follows for the lowest to highest concentrations for each compound: n=27, 29, 29, 34, 25, 34, 35, 25, 32, 29, 31 for PFNA (C); n=23, 34, 37, 29, 40, 31, 16, 23, 30, 38, 16 for PFOA (D); n=44, 35, 44, 32, 51, 50, 47, 40, 38 for PFOS (C); n=32, 45, 38, 43, 40, 41, 33, 15, 11 for PFHxS (F); n=30, 28, 32, 29, 28, 30, 41, 31, 30, 34, 38, 25 for GenX (G); and n=31, 44, 26, 35, 31, 37, 22, 29, 38, 30, 35 for PFBS (H). Data were fitted to a sigmoidal curve using a four-parameter logistic (4PL) model, with X being concentration, through least squares fit.

Following the embryonic exposure, embryos were raised to the juvenile stage for Fishbook testing at 23 dpf. The survival rates of each PFAS treatment condition at 23 dpf are shown in Supplementary Figure 1A-1F. Note that there is a ∼50% attrition during the nursery growth period (5 dpf to 23 dpf) for even the control samples (Supplementary Figure 1A-1F). The range of survival rates for all concentrations of PFNA (41.7%-58.3%; Supplementary Figure 1A), PFOA (26.7%-66.7%; Supplementary Figure 1B), PFOS (53.3%-85%; Supplementary Figure 1C), GenX (41.7%-68.3%; Supplementary Figure 1E), and PFBS (36.7%-73.3%; Supplementary Figure 1F) exposures were each comparable to the survival rates of their respective vehicle control (45% for PFNA, 38.3% for PFOA, 73.3% for PFOS, 50% for GenX, and 51.7% for PFBS). While the range of survival rates for the lower doses of PHFxS (55%-75%; Supplementary Figure 1D) remained comparable to the control (53.3%), the highest two doses of PHFxS (200 µM and 400 µM; Supplementary Figure 1D) appeared to negatively impact survival (25% and 18.3%, respectively). For Fishbook assessment of social behavioral deficits induced by the exposures, we observed a clear dose-response for PFNA (Figure 1C; Supplementary Figure 1G), PFOA (Figure 1D; Supplementary Figure 1H), and PFOS (Figure 1E; Supplementary Figure 1I) but not for PFHxS (Figure 1F; Supplementary Figure 1J), PFBS (Figure 1G: Supplementary Figure 1K), and GenX (Figure 1H) (Supplementary Figure 1L). The IC_50_ values were determined to be 23.85 µM (11.068 ppm) for PFNA (Figure 1C), 40.15 µM (16.63 ppm) for PFOA (Figure 1D), and 0.1892 µM (0.095 ppm) for PFOS (Figure 1E) by fitting the dose-response data to a four-parameter logistic sigmoidal model using nonlinear least-squares regression. These results affirm PFNA and PFOA’s neurodevelopmental toxicity, which are aligned with previous epidemiology findings linking PFNA and PFOA to ASD risk. Our results further demonstrate the toxicity of PFOS in social behavioral development and suggest its potential implication for ASD risk.

### Discovering protein binding targets of PFAS via reverse molecular docking against the human proteome

To discover the protein targets of PFAS, we conducted a reverse molecular docking screen against predicted binding pockets^16^ of the human 3D proteome based on the AlphaFold database^17,18^ (Figure 2A). A single domain was used for pocket prediction for most of the proteins, while multiple domains were used for predictions for a subset of proteins. For each domain, pockets were predicted based on both sequence and 3D structure. The highest-ranking sequence-based and 3D-structure-based pocket predictions were selected for each prot ein domain for the screen. A total of 33446 predicted pockets were screened. Each pocket was screened once via molecular docking for each PFAS. The combined screening results are shown in Supplementary Table 1, screening results for each PFAS are separately shown in Supplementary Tables 2-7.

**Figure 2.**
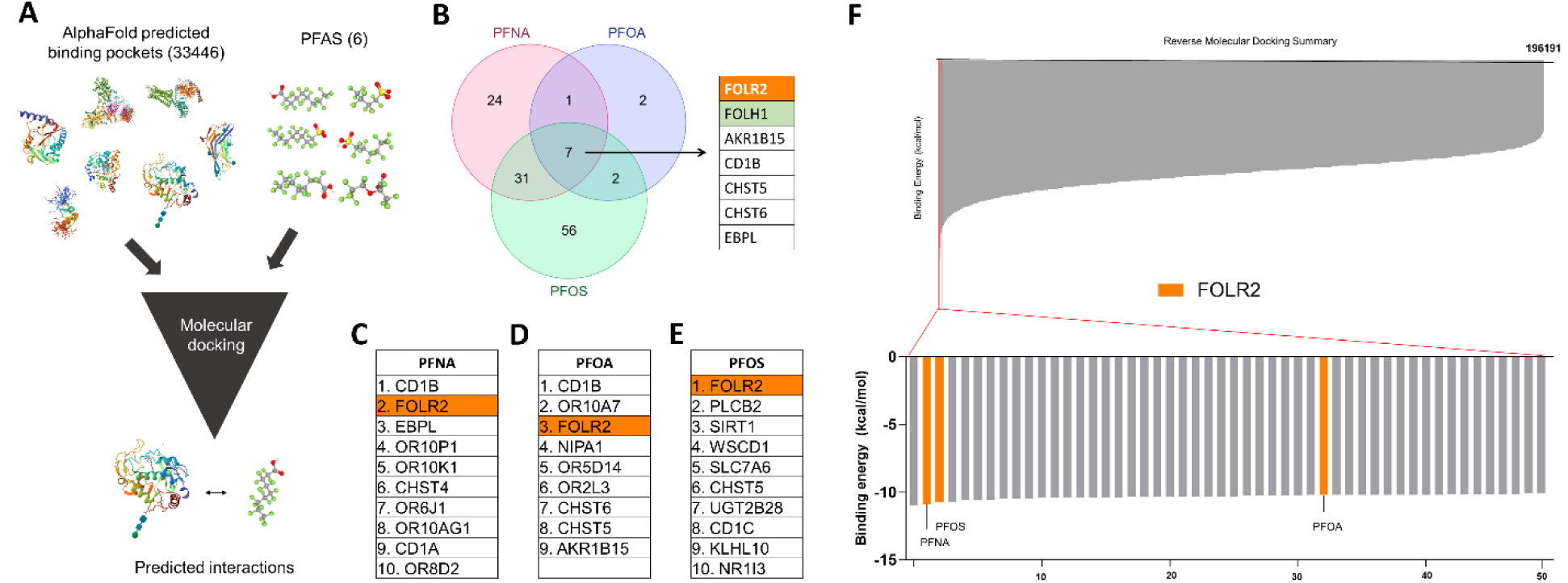
Reverse molecular docking discovered molecular targets of PFAS in the human proteome. (**A**) A schematic drawing demonstrating the reverse molecular docking approach. (**B**) A Venn diagram showing overlaps between the predicted protein targets of PFNA, PFOA, and PFOS. Hits were identified by selecting protein-PFAS interactions with predicted binding energy < −10 kcal/mol. The 7 proteins predicted to bind to all three PFAS are listed in the table pointed by the arrow. (**C-E**) Lists of the top 10 hits with predicted binding energy <-10 kcal/mol for PFNA (C), PFOA (D), and PFOS (E). (**F**) Summary of the reverse molecular docking screen. All molecular docking scores of six EPA-regulated PFAS compounds against predicted binding pockets in 80% of human proteome are shown as a bar chart at the top. The top 50 highest affinity binding interactions (the lowest scores) are zoomed in and shown as a bar chart at the bottom. Interactions between PFAS (PFNA, PFOS, and PFOA) and FOLR2 are highlighted in orange. Ranking is based on average binding scores for each PFAS-domain interaction, calculated from binding energies between each PFAS and binding pockets predicted by structural-based prediction and sequence-based prediction for each protein domain.

We set a cutoff at binding energy of −10 kcal/mol. Using this cutoff, no hit was identified for PFBS and GenX, and only 1 hit was identified for PFHxS (Supplementary Table 5). A number of hits were identified for PFNA, PFOA, and PFOS (Figure 2B & Supplementary Figure 2A). We noticed that folate receptor beta, or FOLR2, consistently appeared among the top hits for the three PFAS (Figure 2B), ranking #2 among the predicted targets of PFNA with a binding energy of −10.9 kcal/mol (Figure 2C), #3 for PFOA at −10.2 kcal/mol (Figure 2D), and #1 for PFOS at −10.73 kcal/mol (Figure 2E). The other two folate receptor subtypes, folate receptor alpha (FOLR1) and folate receptor gamma (FOLR3) were also targeted by PFAS, although at lower affinities with FOLR1 binding to PFNA at an energy of −8.6 kcal/mol, to PFOA at −8.9 kcal/mol, and to PFOS at −9.4 kcal/mol, and with FOLR3 binding to PFNA at an energy of −9.9 kcal/mol, to PFOA at −9.4 kcal/mol, and to PFOS at −10 kcal/mol (Supplementary Tables 2-4). Another two proteins involved in the folate metabolic pathway, folate hydrolase 1 (FOLH1) and methenyltetrahydrofolate synthetase (MTHFS), were also predicted to be bound by PFAS (Figure 2B & Supplementary Tables 2-4), with FOLH1 binding to PFNA at an energy of −10.1 kcal/mol, to PFOA at −10.1 kcal/mol, and to PFOS at −10.1 kcal/mol, and with MTHFS binding to PFNA at −10 kcal/mol, to PFOA at - 9.5 kcal/mol, and to PFOS at −6.5 kcal/mol. The combined outcome of the screening is summarized in Figure 2F and Supplementary Figure 2B.

Consistent with our observations, enrichment analyses identified the folate pathway in PFNA, PFOA, and PFOS targeted proteins among the Gene Ontology (GO) pathways (Figure 3A & Supplementary Figures 3A-3C) and the Kyoto Encyclopedia of Genes and Genomes (KEGG) pathways (Figure 3B & Supplementary Figures 3D-3F). Protein-protein interaction (PPI) analysis using stringDB for the top hits (with predicted binding energy < −10 kcal/mol) of PFNA (Figure 3C), PFOA (Figure 3D), and PFOS (Figure 3E) consistently identified an interaction between FOLR2 and FOLH1. We did not find additional PPIs that consistently connect to FOLR2.

**Figure 3.**
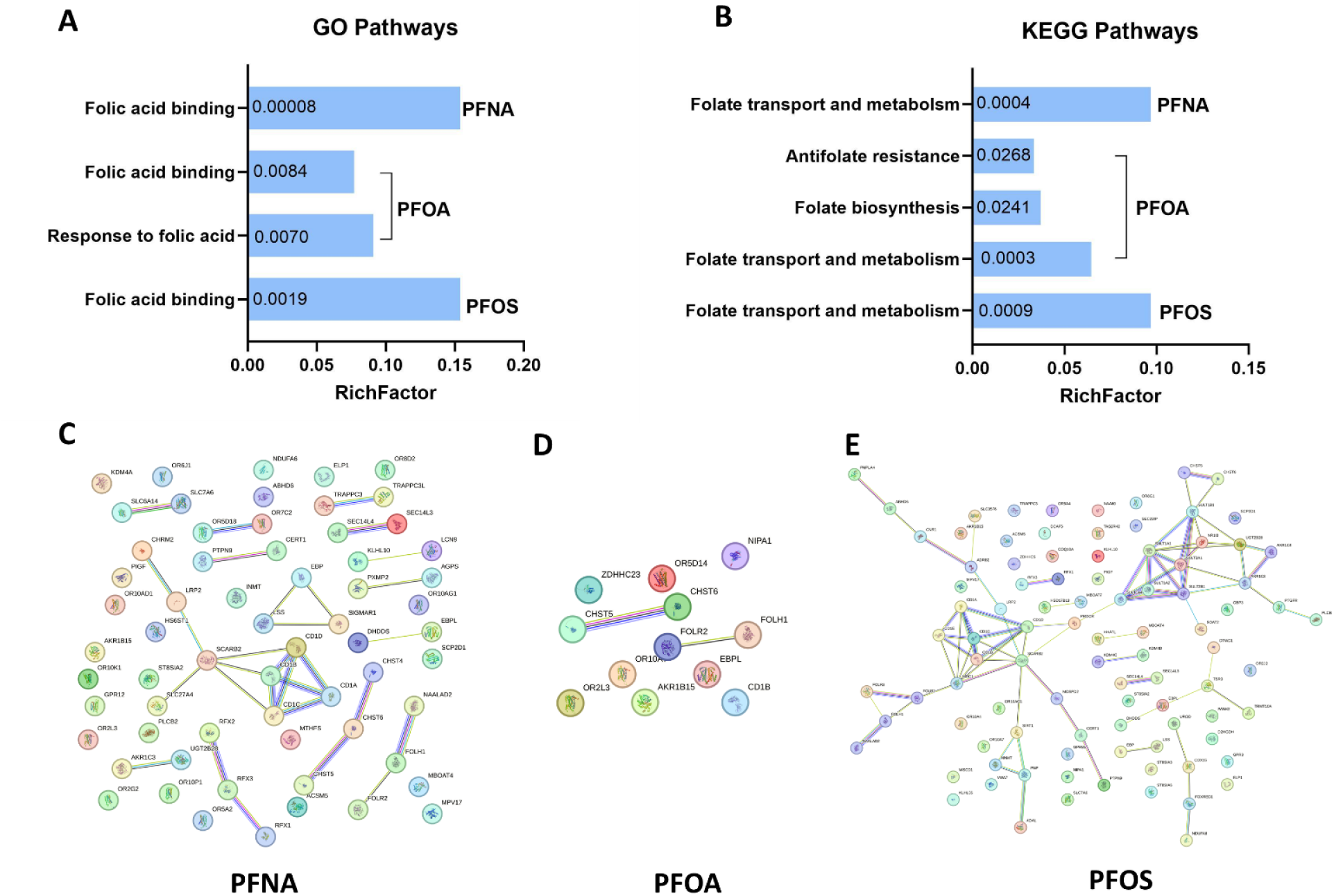
Enrichment analysis and protein-protein interaction analysis consistently identified the folate pathway as a target of PFAS. (**A-B**) Enrichment analyses for the GO (A) and KEGG (B) pathways consistently identified the folate pathway for PFNA, PFOA, and PFOS. Values inside each bar represents adjusted *p*-value. (**C-F**) stringDB analysis for the top hits of PFNA (C), PFOA (D), and PFOS (E) found no consistent PPI networks except a connection between FOLR2 and FOLH1.

### *In silico*, *in vitro*, and *in vivo* validations confirm folate receptor as a biological target of PFAS

To verify FOLR2 as a binding target for PFNA, PFOA, and PFOS, and compare their binding affinities with the endogenous ligand of FOLR2, folic acid (or folate), in its natural binding pocket, we conducted molecular docking using a 3D structure of FOLR2 from the Protein Data Bank (PDB) in complex with folate (4KMZ). We found that the natural binding pocket of folate (Figure 4A) can be effectively bound by PFNA (Figure 4B), PFOA (Figure 4C), and PFOS (Figures 4D). In fact, an average of 5 random docking trials showed that PFNA, PFOA, and PFOS all bind to FOLR2 at a higher potency (−11.04±0.2191 kcal/mol, −10.34±0.1949 kcal/mol, and −10.28±0.04472 kcal/mol, respectively) compared to folate (−9.68±0.04472 kcal/mol), while PFHxS, PFBS, and GenX bind to FOLR2 at a lower potency compared to folate (−9.3±0.0 kcal/mol, −8.82±0.2049 kcal/mol, and - 8.1±0.0 kcal/mol, respectively) (Figure 4E). We also conducted molecular docking with FOLR1 using a 3D structure of FOLR1 from the PDB in complex with folate (4LRH) (Supplementary Figures 4A-4D) and found a similar pattern of differential binding affinity among the PFAS species: PFOS and PFNA bind to FOLR1 at a potency higher than folate (PFOS: −10.08±0.08357 kcal/mol, PFNA: −9.8±0.2236 kcal/mol, folate: −9.58±0.1095 kcal/mol), whereas PFOA, PFHxS, PFBS, and GenX bind to FOLR1 at a lower potency compared to folate (−9.3±0.0 kcal/mol, −8.7±0.0 kcal/mol, −8.4±0.0 kcal/mol, and −7.7±0.0 kcal/mol, respectively) (Supplementary Figure 4E). Consistent with a previous report which associated PFAS exposure to lower folate levels in human blood^23^, our findings suggest that exposure to PFNA, PFOA, and PFOS may out-compete FOLR2’s natural ligand, folate, and lead to reduced folate uptake *in vivo*.

**Figure 4.**
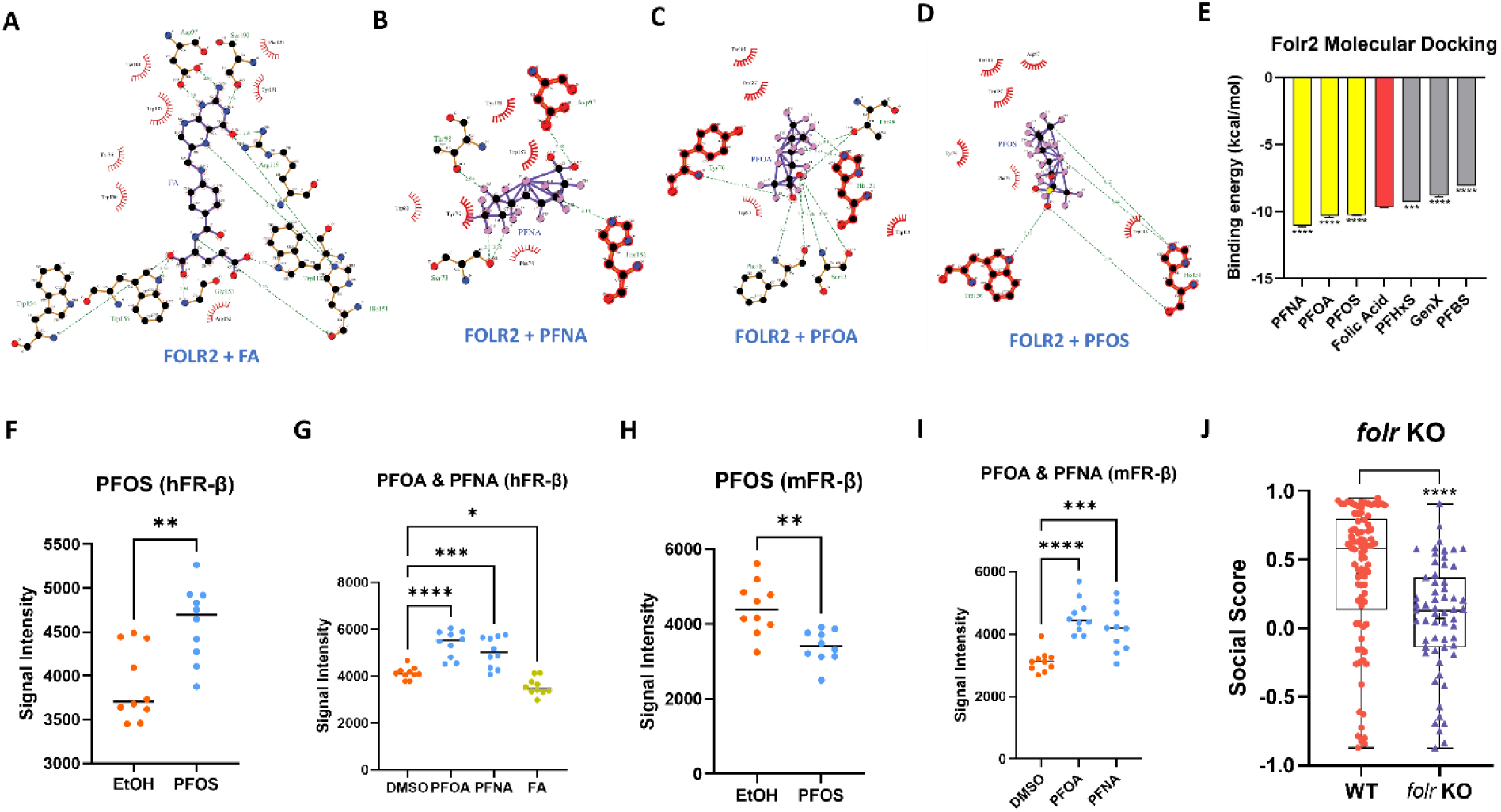
*In silico*, *in vitro*, and *in vivo* assays validated folic acid receptor as the biological target of PFAS. (**A-D**) *In silico* validation via molecular docking. Showing docking images for FOLR2 against folic acid (FA) (A), PFNA (B), PFOA (C), and PFOS (D). FOLR2 structure was acquired using the PDB identifier 4KMZ. (**E**) Comparing the binding energies for folic acid and the six EPA-regulated PFAS against FOLR2. Binding energies are shown as average and standard deviation of 5 independent docking attempts. Significance values are calculated to compare PFAS data with folic acid. ***: *p* < 0.001, ****: *p* < 0.0001. (**F-G**) Dot blot analyses detected significantly different protein levels in the pooled soluble fractions of PISA samples between PFAS-treated samples and vehicle controls. PISA was conducted using recombinant human FR-β protein (hFR-β) against PFOS (10 µM) (F), PFOA (1600 µM) (G), and PFNA (400 µM) (G). Folic acid (12.5 µM) (FA) was used as a positive control (G). Datapoints represent raw signal intensities of the dot blot assays. Each sample were tested at n=10 replicates. (**H-I**) PISA and dot blot analyses revealed similar results using recombinant mouse FR-β protein (mFR-β) against PFOS (H), PFOA (I), and PFNA (I). Datapoints represent raw signal intensities of the dot blot assays. Each sample were tested at n=10 replicates. (**J**) CRISPR-Cas9 induced F0 knockout of the zebrafish folic acid receptor *folr* induced social deficits in zebrafish. Each datapoint represents the social score of an individual fish. A total of n=86 wild-type control fish (WT) and n=59 *folr* KO fish were tested. Significance was calculated by one-way ANOVA and Dunnett’s multiple comparison test when comparing multiple groups (E, G, I) or two-tailed Student’s *t* test when comparing two groups (F, H, J). *: *p* < 0.05, **: *p* < 0.01, ***: *p* < 0.001, ****: *p* < 0.0001.

We then examined whether PFNA, PFOA, and PFOS can physically bind to the FR-β protein. We applied an *in vitro* assay called proteome integral solubility alteration (PISA), which detects the binding of a chemical to its target protein by comparing the soluble portion of the compound-protein mixture with a control mixture following a temperature gradient treatment^24^. His-tagged recombinant human and mouse FR-β proteins were incubated with PFOS, PFOA, PFNA, folic acid, or vehicle control for 30 minutes at room temperature. The treated samples were subjected to a thermal gradient (34–66°C) in a thermocycler, pooled, and centrifuged to isolate soluble proteins. Supernatants were applied to a nitrocellulose membrane at n=10 replicates for dot blot analysis. We found that PFOS (Figure 4F), PFOA, PFNA, and folic acid (Figure 4G) all significantly altered the amount of protein remaining in the soluble fraction after thermal gradient treatment, demonstrating their bindings with the human FR-β protein. Similar results were observed for PFOS (Figure 4H), PFOA, and PFNA (Figure 4I) against the mouse FR-β protein. These results demonstrate that PFOS, PFOA, and PFNA can indeed physically bind to the FR-β protein.

Finally, we investigated whether folate deficiency contributes to social behavioral deficits in our zebrafish social behavior model. The zebrafish only possess one gene encoding the folate receptor, *folr*. To evaluate how *folr* loss-of-function affects sociality, we knocked out the zebrafish *folr* using CRISPR/Cas9 by injecting fertilized eggs with a mixture of three *folr* gRNAs and Cas9 protein. The resulting F0 mutants show significant social deficits compared to wild-type controls (*p*<0.0001, Cohen’s *d*=0.70 indicating a medium effect) (Figure 4J), demonstrating that folate pathway is required for social behavioral development in zebrafish. The F0 CRISPR/Cas9 knockout strategy has been shown to faithfully reproduce disease-relevant phenotypes without the confounding influence of compensatory gene expression^25,26^: by simultaneously targeting multiple loci within a single gene, this approach has been reported to consistently achieve over 90% biallelic knockout efficiency and enable rapid phenotypic assessment in F0 animals^26–28^. Indeed, PCR amplification of the gRNA-targeted genomic regions using genomic DNA extracted from injected F0 embryos show that all three gRNAs ablated the WT amplicon, indicating that all three gNRAs effectively mutated the targeted *folr* gene (Supplementary Figure 5).

### Folic acid rescued social deficits induced by PFAS through embryonic co-exposure

Finally, to evaluate whether PFAS induces social deficit through inhibiting the folate pathway, we attempted to rescue social deficits caused by PFAS exposure by co-exposing with folic acid (Figure 5A). We hypothesized that certain doses of folic acid may outcompete PFAS binding, restoring folate metabolism in the developing embryos, without causing developmental toxicity. We co-exposed the three PFAS species including 400 µM PFNA, 1600 µM PFOA, and 10 µM PFOS with a range of folic acid doses (7 doses ranging from 3.125 µM to 200 µM in 2-fold increments) during the first 3 days of embryonic development. Folic acid co-exposure effectively rescued social deficits in all three PFAS treatments. PFNA was rescued by folic acid at the lowest applied doses of 3.125 µM (*p*=0.008, Cohen’s *d*=0.74 indicating a medium effect) and 25 µM (*p*=0.03, *d*=0.58 indicating a medium effect) (Figure 5B & Supplementary Figure 6A). Both PFOA and PFOS showed a bell-shaped rescue curve when co-exposed to folic acid. PFOA was most effectively rescued by folic acid at 6.25 µM (*p*=0.018, *d*=0.88 indicating a large effect) to 12.5 µM (*p*=0.0095, *d*=0.95 indicating a large effect) (Figure 5C & Supplementary Figure 6B), whereas PFOS was robustly rescued by 12.5 µM (*p*=0.0002, *d*=1.16 indicating a large effect), 25 µM (*p* <0.0001, *d*=1.39 indicating a large effect), 50 µM (*p*=0.02, *d*=0.82 indicating a large effect), and 100 µM (*p*=0.017, *d*=0.77 indicating a medium effect) folic acid treatments (Figure 5D & Supplementary Figure 6C). These results indicate that PFAS likely induces social deficit by inhibiting the folate pathway, and this inhibition can be prevented by embryonic co-exposure to folic acid.

**Figure 5.**
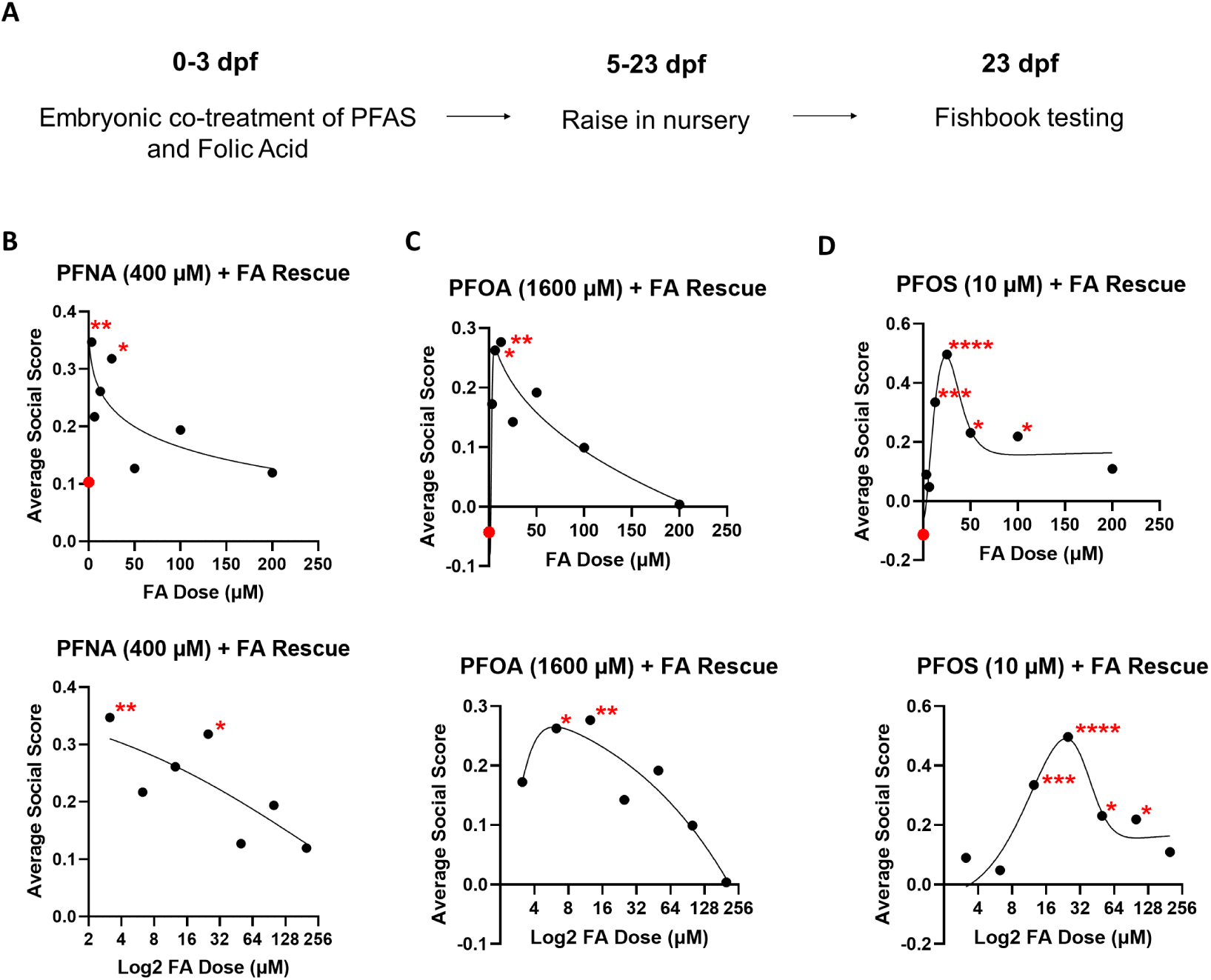
**Embryonic co-exposure to folic acid rescued social deficits induced by PFAS.**(**A**) A schematic ofthe co-treatment and testing workflow. (**B-D**) Dosage curves generated by plotting average social scores (y-axis)against folic acid (FA) concentrations (x-axis). Showing both linear (top)and log2 (bottom) scales for the x-axis.All treatment groups in each experiment were exposed to 400 μM PFNA (B),1600 μM PFOA (C),or10 μM PFOS (D),respectively, plus a range of folic acid (FA) doses (3.125 μM to 200 μM) to assess its rescue effect.Red dotsmarks the no-FAcontrol(0 μMFA)ofeach experimentin the linearplots.Sample sizesare asfollowsforthe lowestto highest FA concentrations in each rescue experiment:n=43, 44, 42, 46, 42, 29, 45, 22for PFNA+FA (B); n=28, 29, 29, 32, 29, 30, 27, 18forPFOA+FA(C);and n=32, 23, 36, 37, 20, 25, 31, 40for PFOS+FA(D). Data werefitted toa bell-shaped curve using a four-parameter logistic(4PL)model,with Xbeing concentration,through leastsquaresfit. Significance was calculated by one-wayANOVA and Dunnett’s multiple comparison test.*:*p*< 0.05, **:*p*< 0.01,***:*p*< 0.001,****:*p*< 0.0001.

## DISCUSSION

Our study provides critical insights into the neurodevelopmental toxicity of six EPA-regulated PFAS compounds—PFNA, PFOA, PFOS, PFHxS, PFBS, and GenX—on social behavioral development in zebrafish. Using the Fishbook assay, we observed distinct dose-dependent responses for PFNA, PFOA, and PFOS, which displayed neurodevelopmental impacts on social behavior in the exposed embryos, while PFHxS, PFBS, and GenX exhibited no apparent dose-dependent effects on social behavior within the tested concentration range. IC_50_ values were established for PFNA, PFOA, and PFOS, reinforcing epidemiological links between PFNA, PFOA, and ASD risk, while suggesting a potential contribution of PFOS to ASD risk. This specificity in behavioral effects aligns with our molecular docking results, where PFNA, PFOA, and PFOS displayed stronger binding affinities to the folate receptor FOLR2, suggesting that these PFAS may disrupt folate metabolism and thereby influence neurodevelopmental pathways implicated in social behavioral development. This proposed mechanism is consistent with previous human biomonitoring evidence showing an inverse association between folic acid and PFAS levels in human red blood cells and serum samples^23^, indicating that PFAS exposure in humans likely impedes folic acid uptake or bioavailability. Based on these convergent behavioral, molecular docking, and epidemiological observations, we hypothesize that a subset of PFAS with specific physicochemical properties disrupt neurodevelopment by interfering with folate receptor-mediated folate transport during critical developmental windows, thereby increasing susceptibility to ASD-relevant social behavioral deficits.

PFAS exposure is nearly ubiquitous for people living in the U.S., but the magnitude and composition of exposure vary substantially across different exposure scenarios such as the general population, communities at risk of exposure, and occupational exposure settings. Understanding these real-world exposure levels is essential for interpreting our results and evaluating the translational relevance of our findings. According to the National Health and Nutrition Examination Survey (NHANES), in the U.S. general population (GP), median serum concentrations of PFOS and PFOA in women of childbearing age have declined significantly over the past two decades due to regulatory actions and changes in industrial practices. Among the PFAS species examined in this study, PFOS decreased from approximately 24 ng/mL (48 nM) in 1999–2000 to 3 ng/mL (6 nM) in 2017–2018, and PFOA decreased from 5 ng/mL (12 nM) to 1 ng/mL (2.4 nM) over the same period. In contrast, PFNA exhibited an initial rise from 0.5 ng/mL in 1999–2000 to 1.0 ng/mL (2 nM) in 2009–2010 before declining to 0.3 ng/mL (0.5 nM) in 2017–2018. Despite these downward trends, PFAS remain detectable in the blood of more than 97% of Americans, reflecting their environmental persistence, bioaccumulation, and widespread distribution in food and drinking water. Exposed Communities (EC) near industrial plants, military installations, or municipal water supplies continue to experience disproportionately high PFAS exposure. For example, residents served by the Little Hocking Water Association in Ohio had a mean PFOA serum level of 227.6 ng/mL (0.55 µM), while exposed communities in Decatur, Alabama reported a PFOS concentration of 39.8 ng/mL (80 nM). These levels are tens to hundreds of times higher than those observed in the general population. Occupational exposure (OE) settings represent the highest end of the PFAS exposure spectrum. Historical data from PFAS manufacturing workers in Decatur, Alabama, showed mean serum concentrations approaching 899 ng/mL (2.2 µM) for PFOA and 941 ng/mL (1.9 µM) for PFOS^29^, which are orders of magnitude above exposure levels of the general population (**Table 1**).

**Table 1.** Human exposure levels of PFOS, PFOA, and PFNA.

|  | **General Population Low (GPL)** | **General Population High (GPH)** | **Exposed Communities (EC)** | **Occupational Exposure (OE)** |
| --- | --- | --- | --- | --- |
| **PFOS** | 3 ng/mL<br>(6 nM) | 24 ng/mL<br>(48 nM) | 39.8 ng/mL<br>(80 nM) | 941 ng/mL<br>(1.9 µM) |
| **PFOA** | 1 ng/mL<br>(2.4 nM) | 5 ng/mL<br>(12 nM) | 227.6 ng/mL<br>(0.55 µM) | 899 ng/mL<br>(2.2 µM) |
| **PFNA** | 0.3 ng/mL<br>(0.5 nM) | 1.0 ng/mL<br>(2 nM) | N.A. | N.A. |

Our preliminary data revealed an IC_50_ value of 189.2 nM for PFOS (Figure 1E), indicating that human PFOS exposures (GP: 6-48 nM, EC: 80 nM, OE: 1.9 µM) may approach an at-risk range. Moreover, because PFOS, PFOA, and PFNA share a conserved molecular target, their combined presence in humans may interfere with folate transport during neurodevelopment through additive or synergistic effects, even though individual serum levels are lower than their respective toxic doses in zebrafish. This possibility raises the need to consider cumulative PFAS burden rather than focusing on single compounds. In short, among the six PFAS species regulated by the EPA, PFOS remains the most relevant to ongoing human exposure, yet PFOA and PFNA cannot be overlooked given their shared folate receptor-binding potential and likelihood of additive neurotoxicity. Investigating these compounds and other environmental PFAS species with folate receptor-binding activities is therefore crucial for exposure mitigation, mechanistic understanding, and health-risk assessment in vulnerable populations.

The selective behavioral and molecular effects observed among the tested PFAS suggest a potential structure-activity relationship. PFNA, PFOA, and PFOS share longer carbon chain lengths and greater hydrophobicity compared to PFBS, PFHxS, and GenX, properties that may enhance protein binding affinity, membrane interactions, and bioaccumulation. These physicochemical features could facilitate stronger or more sustained interactions with folate receptors or associated transport machinery, contributing to their observed neurodevelopmental effects. However, additional factors such as functional head groups, branching, and differential tissue distribution may also play important roles and warrant systematic investigation in the future.

Recent studies have revealed a potential role of folic acid in ASD^22^. Low maternal folate levels have been associated with a higher incidence of ASD in offspring^30–32^. A number of studies have linked maternal folic acid supplementation with reduced risk of ASD in offspring^32–38^. For example, maternal folic acid supplementation at 600 µg has been associated with a significantly reduced risk of ASD in offspring^33^, suggesting that adequate folate levels during pregnancy may be beneficial for disease prevention. Specifically to PFAS exposure, a recent epidemiology study based on the Shanghai Birth Cohort detected an association between prenatal PFOA exposure and increased risk of autistic traits in the offspring, and found that the PFOA-induced autistic risk was mitigated by pre-pregnancy folic acid supplementation^11^, directly supporting our findings. However, the dosage of folic acid supplementation appears to affect its protective effects, with higher or lower levels of supplementation associated with increased risk of ASD^35^. Supplementation with folic acid and/or folinic acid in children diagnosed with ASD improved their neurological and behavioral symptoms^39–43^. Several studies have reported an elevated prevalence of serum auto-antibodies against folate receptor alpha in children with ASD^44–46^. Finally, polymorphisms in the methylenetetrahydrofolate reductase (MTHFR) gene have been associated with ASD risk^47–53^. These findings align with our observations that PFAS-induced folate pathway disruption can lead to social behavioral deficits.

The folate pathway may contribute to ASD through several potential mechanisms. Folic acid is required for DNA methylation, a critical process during brain development: it is a source of the one-carbon group used to methylate DNA and required for the production of S-adenosylmethionine, the primary methyl group donor for DNA methylation. Folic acid is also essential for neural tube formation. It is worth noting that the optimal protective effects of folic acid against ASD was observed when folic acid was taken during preconception and the first trimester of pregnancy^33,38^, the latter of which overlaps with the period of neural tube formation. Both DNA methylation and neural tube formation are critical processes for brain development, pointing to potential mechanisms of action for folic acid’s involvement in ASD. Additionally, folic acid is essential for the production of neurotransmitters that affect mood, motivation, stress, and cognitive performance, including serotonin, norepinephrine, and dopamine^54,55^, influencing neural signaling pathways that are often disrupted in ASD.

Some studies indicate a possible link between elevated maternal folate levels and an increased risk of ASD, although findings are inconsistent and complex due to variations in individual metabolism, timing, and dosage of supplementation^56^. These findings suggest that while folic acid is crucial for brain development^19–21^, there is a need for more nuanced guidance on optimal intake levels to balance benefits with potential risks, especially for populations at risk for ASD. This is aligned with our finding that while moderate to low doses of folic acid exerted beneficial rescue effects on social behavior when co-exposed to PFAS, higher doses showed significantly reduced rescue potencies.

Although prior work has utilized reverse molecular docking on selected proteins to predict binding targets of the environmental toxicant toluene^57^, to our knowledge, this work represents the first proteome-wide reverse molecular docking in toxicology to discover *in vivo* protein targets for environmental toxicants. In reverse molecular docking, a small molecule drug or toxicant is virtually docked into the potential binding pockets of a large set of proteins. The binding characteristics of these interactions are then analyzed to rank the protein targets based on their binding affinities to the small molecules. This approach is different from traditional molecular docking which typically screens a large number of small molecules against a predetermined protein target, with the goal of identifying a ligand with high binding affinity to this protein. The successful application of proteome-wide reverse molecular docking in toxicology demonstrates a rapid and systematic *in silico* approach to predict protein targets for key environmental toxicants, which is expected to significantly promote the advance of mechanistic toxicology.

While our findings support a folate receptor-centered mechanism, alternative and complementary pathways may also contribute to PFAS-induced neurodevelopmental toxicity. PFAS are known to interact with multiple biological targets, including nuclear receptors such as PPARs^58–61^, lipid metabolism pathways^62–66^, thyroid hormone signaling^67–69^, oxidative stress responses^70–72^, and calcium homeostasis^73–75^, all of which can influence brain development and behavior. Disruption of these pathways could act independently of, or synergistically with, folate pathway perturbations to shape neurodevelopmental outcomes. In addition, several limitations of the present study should be acknowledged. Although zebrafish provides a powerful vertebrate model for high-throughput behavioral and mechanistic studies, differences in PFAS toxicokinetics, placental transfer, and folate transport mechanisms between zebrafish and humans may influence dose-response relationships and translational interpretation. Moreover, our molecular docking analyses predict binding affinity but do not directly measure *in vivo* receptor occupancy or functional inhibition of folate transport, which will require future biochemical and genetic validation.

In summary, this study provides evidence that specific PFAS, notably PFNA, PFOA, and PFOS, may contribute to social behavioral deficits through their interaction with folate receptors, an effect mitigated by co-exposure to folic acid. Our findings align with existing evidence on the association between folate pathway disruptions and ASD risk, contributing to a deeper understanding of the biochemical mechanisms by which environmental toxicants may influence neurodevelopmental and disease outcomes. Furthermore, the ability of folic acid to rescue PFAS-induced deficits highlights the potential of folate supplementation to counteract PFAS toxicity in developing embryos. Together, these results underscore the need for further research into folate-based interventions and the long-term implications of exposure to various species of PFAS on neurodevelopmental disorders, particularly ASD. In addition, by incorporating the innovative proteome-wide reverse molecular docking approach, we provide a comprehensive framework for advancing the field of mechanistic toxicology through the identification of novel toxicological targets.

## MATERIALS AND METHODS

### Zebrafish husbandry

Zebrafish were housed at 26°C–27°C on a 14-hour light, 10-hour dark light cycle. Wild-type AB strain was used for all experiments. Nursery fish were fed twice a day at roughly 9 AM to 12 PM and 2 PM to 5 PM during the weekdays and once during weekends at no set time. Nursery fish were fed only rotifers from 5 dpf to 10 dpf, and were fed a mixture of rotifers and artemia between 10 dpf to 23 dpf. All zebrafish experiments were approved by the Institutional Animal Care and Use Committee at the University of Washington.

### Chemical exposure

Fertilized embryos were sorted and transferred into 60 mm diameter and 20 mm deep glass petri dishes at 30 embryos per dish. Each dish was filled with 10 mL of HEPES-buffered E3 medium. Compound stocks were prepared in their specified solvents to maintain stability and stored at - 20°C. Each compound was added to duplicate dishes (30 embryos per dish, two dishes per condition) at various concentrations, with negative controls receiving an equal volume of the corresponding solvent. Embryos were exposed without dechorionation. To prevent media contamination, dead embryos were removed at 1 and 2 days post-fertilization (dpf). At 3 dpf, all viable larvae were rinsed with E3 medium and transferred into clean petri dishes containing fresh E3 medium. At 5 dpf, larvae from each petri dish were transferred to separate nursery tanks at a final density of no more than 50 fish per tank and raised to 23 dpf for Fishbook assays.

The following per- and polyfluoroalkyl substances (PFAS) were obtained from Cayman Chemical (Ann Arbor, MI, USA): Perfluorooctanesulfonic Acid (PFOS, CAS# 1763-23-1, Item no. 37233), Perfluorooctanoic Acid (PFOA, CAS# 335-67-1, Item no. 37232), Perfluorononanoic Acid (PFNA, CAS# 375-95-1, Item no. 37250), Perfluorohexanesulfonic Acid (PFHxS, CAS# 355-46-4, Item no. 37248), Perfluorobutanesulfonic Acid (PFBS, CAS# 375-73-5, Item no. 37243), and Folic Acid (CAS# 59-30-3, Item no. 20515). Ammonium perfluoro(2-methyl-3-oxahexanoate) (GenX, CAS# 62037-80-3, Item no. C23990) was obtained from Sigma-Aldrich. PFOS came as a solution in 100% ethanol. PFBS came in liquid form. To prepare stock solutions, PFOA, PFNA, and folic acid were dissolved in HEPES-buffered E3 media, GenX and PFHxS were dissolved in dimethyl sulfoxide (DMSO).

### Fishbook assay

The Fishbook assay was run as described previously^13^. Briefly, the test arena consists of a total of 44 3D printed, 10-mm-deep, 8.5-mm-wide, and 80-mm-long rectangular chambers grouped together in parallel with each other. Each chamber is divided into three compartments by two transparent acrylic windows (1.5 mm thick): a 60-mm-long middle testing chamber to place the test subject and two 8.5-mm-long end chambers to place the social stimulus fish or remain empty, respectively. Test subjects (PFAS treated 23 dpf fish or controls) were each placed inside an individual test chamber using a plastic transfer pipette with its tip cut off to widen the opening. A 3D printed white comb-like structure was placed in front of the social stimulus compartment to block the test subject’s visual access to social stimulus fish before testing begins. After test subjects were placed inside the chambers, the arena was placed inside the imaging station, and the combs were removed to visually expose the social stimulus fish to the test subjects. Following a brief acclimation period, a 10-min test session was video recorded.

Videos were streamed through the software Bonsai^76^. Videos were analyzed in real time during recording, and the frame-by-frame x and y coordinates of each fish relative to its own test compartment were exported as a CSV file. Data were analyzed using custom Python scripts to calculate social scores and generate tracking plots. Social score was defined as a fish’s average y-axis position for all frames. The middle of each test chamber was designated as the origin of the y axis, with an assigned value of zero. A value of 1 was assigned to the end of the chamber next to the social stimulus fish and a value of −1 to the other end of the chamber next to the empty control compartment. In this coordinate system, all social scores have values between −1 and 1. A higher social score demonstrates a shorter average distance between a test subject and a social stimulus fish during a test, which suggests a stronger social preference.

### CRISPR-Cas9 gene knockout

We designed 3 sets of sgRNAs targeting the zebrafish *folr* gene to maximize efficiency of F0 knockout^26–28^. CHOPCHOP^77^ was used for the sgRNA design. The *folr* sgRNA sequences designed are as follows: gRNA1 “ATTTAGGTGACACTATATGAGGGACAGTTATATCAGCGTTTTAGAGCTAGAAATAGCAAG”, gRNA2 “ATTTAGGTGACACTATATGTATCCAGGGCCCCAGGTGGTTTTAGAGCTAGAAATAGCAAG”, and gRNA3

“ATTTAGGTGACACTATAGTCCATGTTCTGCTCAGGTGGTTTTAGAGCTAGAAATAGCAAG”. sgRNAs targeting early coding exons were selected to introduce nonsense mutations as early as possible and increase likelihood of loss-of-function mutations. The SP6 promoter sequence was added before the target RNA sequence, followed by an overlap adapter, which is complementary to the 5’ end of an 80 bp constant oligo according to the published protocol^78^. Oligonucleotides were synthesized by Eurofins Scientific. Double-stranded DNAs were generated by Phusion Hot-Start Flex DNA Polymerase (New England Biolabs) using the gene specific oligos and the constant oligo. The double-stranded DNA was cleaned up using the Zymo Clean and Concentrator-5 kit (Zymo Research). *In vitro* sgRNA transcription was conducted using the MEGAscript SP6 transcription kit (Thermo Fisher Scientific) and cleaned up using the Zymo RNA Clean and Concentrator-5 kit (Zymo Research).

To perform zebrafish embryonic microinjections, pairs of male and female adult zebrafish were kept in a mating cage overnight while separated by a divider. 1-cell-stage embryos were collected the next day early in the morning immediately before injection by pulling out the divider to allow breeding. Injection solution was prepared by combining the three sgRNAs together with 2 µM Spy Cas9 NLS and 1X NEBuffer (New England Biolabs). All three gRNAs were injected together to induce multiple point mutations in the same gene. Approximately 1 nL of sgRNA–Cas9 RNP complex was injected into each 1-cell-stage embryo using a pneumatic microinjector (Parker Instrumentation). Approximately 300 embryos were injected with an estimated 60% survival rate 24 hours post-injection. To verify successful knockout of *folr*, we extracted genomic DNA from injected embryos and amplified the gRNA-targeted genomic regions using PCR. The following forward and reverse primers were used: sgRNA1 forward primer: GTGTGATGTGTCTGTGTTGTGC; sgRNA1 reverse primer: TGTGTGTTGCAGTGAAGTGTGT; sgRNA2 forward primer: AAGAAGCGCATCAGGATAACTC; sgRNA2 reverse primer: CTGATGTTTAAAACTGGTGGCA; sgRNA3 forward primer: CAACGTGATTGCCTCTATGAAA; sgRNA3 reverse primer: AAACTCAGAAGCATGTGGTGTG. PCR was conducted in a Bio-Rad thermal cycler using a 20µL reaction volume containing 10µL of Taq 2X Master Mix (NEB) and 0.4µL of each primer. The following program was used: a 3 minute initial denaturation at 95°C, followed by 39 cycles of 15 sec at 95°C for denaturation, 30 sec at 54 °C for annealing, and 30 sec at 68 °C for extension, and a final extension step at 68 °C for 5 min. PCR products were examined by electrophoresis at 100 V for 30 minutes in a 1.5% (w/v) agarose gel in 1 x TAE buffer. The gel was imaged on a UV light box at 365 nm. Lack of amplification products or smeared amplification products indicate successful knockout (Supplementary Figure 5).

### Reverse molecular docking screen and molecular docking validation

All *in silico* calculations were performed using a Digital Storm workstation with AMD Ryzen Threadripper PRO 5955WX (16-Core) processor (Up to 4.0 GHz), 128GB DDR4 memory, and NVIDIA RTX A2000 6GB graphics card. Both reverse molecular docking screen and molecular docking validation were conducted using Autodock Vina^79^ and Open Babel^80^. Ligand structures were downloaded from PubChem as sdf files and converted to pdbqt. Binding pocket predictions were acquired from the HPRoteome-BSite database^16^ which covers over 80% of the UniProt entries in the AlphaFold human proteome database^17,18^. In the HPRoteome-BSite database, a binding pocket was predicted for a single domain for most of the proteins, whereas a subset of proteins had a binding pocket predicted for multiple domains. For each domain, pockets were predicted based on either the peptide sequence or the 3D structure of the domain; both types of predictions were used for the screen. The highest-ranking sequence-based predictions and the highest-ranking 3D-structure-based pocket predictions were respectively selected for each protein domain, resulting in a total of 33446 predicted pockets selected for the screen. The domain structures were downloaded from the HPRoteome-BSite database and prepared for molecular docking by removing all water molecules, ions, and other substructures. The docking coordinates of each predicted binding pocket were provided by the HPRoteome-BSite database. Pocket size of 60 × 60 × 60, energy range of 3, number of modes of 9, and an exhaustiveness of 8 were employed for the molecular docking screen and validation. Each pocket was screened once for each PFAS.

### Protein-protein interaction analysis

Protein-protein interaction (PPI) analysis was conducted using STRING^81^ (https://string-db.org/).

### Gene enrichment analysis

Molecular targets with a binding energy of less than −10 kcal/mol from the reverse molecular binding virtual screen were selected for gene enrichment analysis. Gene Ontology (GO), and Kyoto Encyclopedia of Genes and Genomes (KEGG) enrichment analysis were conducted using the clusterProfiler package in R (version 4.12.5). The GO analysis included Biological Process (BP), Cellular Component (CC), and Molecular Function (MF) categories, with the org.Hs.eg.db database used for annotation (v3.19.1). Results were ranked based on RichFactor and adjusted *p*-value. Adjusted *p*-value was calculated using the Benjamini-Hochberg (BH) method with a cutoff of 0.05 for significance. RichFactor is defined as the ratio of input genes that are annotated under a certain term to all genes annotated under this term. Results were visualized as bar plots using the ggplot2 package (v3.5.1).

### Proteome integral solubility alteration (PISA) assay

His-tagged recombinant human folate receptor beta (FR-β) protein (ProSpec, PRO-2673, amino acids 22-230) and mouse FR-β protein (Antibodies-Online, ABIN7274670, amino acids 21-227) were reconstituted in distilled water at a concentration of 500 µg/mL. The proteins were then diluted to 50 µg/mL and incubated with 10 µM PFOS, 400 µM PFNA, 1600 µM PFOA, 12.5 µM folic acid, or an equal volume of vehicle control (DMSO or ethanol) for 30 minutes at room temperature on a rotator. All compounds were prepared as 100x stock solutions compared to their final concentration. Each compound-treated sample was aliquoted into 9 PCR tubes and subjected to a 5 min temperature gradient treatment (from 34°C to 66°C, in 4°C increments) using a Bio-Rad Thermocycler. After temperature treatment, the samples were placed on ice for at least 5 minutes. Contents of the 9 PCR tubes were pooled into a new 1.5 mL tube and centrifuged at 15,000g for 30 minutes at 4°C. The supernatant was transferred to a new 1.5 mL tube. For dot blot analysis, 2 µL of supernatant was applied onto a nitrocellulose membrane, and the procedure was repeated 10 times for each sample (10 technical replicates). Each membrane was air-dried and blocked for 30 minutes in Tris-buffered saline with Tween 20 (TBST) containing 2% bovine serum albumin (BSA). The membrane was then incubated with an anti-His Tag Antibody (SinoBiological, 105327-MM02T) diluted 1:1000 in TBST containing 0.1% BSA and 0.1% sodium azide for 30 minutes. Following primary antibody incubation, the membrane was washed 3 × 5 min by TBST, and incubated with horseradish peroxidase (HRP)-conjugated secondary antibody (Jackson ImmunoResearch) diluted 1:10,000 in TBST containing 0.1% BSA for 30 minutes. After washing in TBST once for 15 minutes and twice for 5 minutes, the membrane was washed once in TBS for 5 minutes, and stored in TBS at 4°C if needed. For imaging, each membrane was exposed to ECL substrate (Bio-Rad) and scanned using the iBright Imaging System (Invitrogen) with chemiluminescence smart exposure. Quantification of the dot blot image was performed using FIJI^82^.

### Statistical analysis

Graphs were generated using GraphPad Prism. Two-tailed Student’s *t* test was used to analyze data when there were two groups. For analysis of multiple groups, normal distribution of datasets was confirmed using Shapiro-Wilk test, and if more than half the data was normally distributed and standard deviations fell within the variance ratio, one-way analysis of variance (ANOVA) assuming gaussian distribution and equal standard deviation was performed, using Dunnett’s multiple comparison test. *P* values less than 0.05 were considered significant.

## ACKNOWLEDGEMENTS

We thank the University of Washington Office of Comparative Medicine for providing zebrafish husbandry support. This work was supported by the National Institute of Environmental Health Sciences (NIEHS) of the NIH under the award number R00ES031050. The content is solely the responsibility of the authors and does not necessarily represent the official views of the NIH.

## AUTHOR CONTRIBUTIONS

M.J. and A.X.K. designed and conducted the experiments and analyzed data. A.E. and K.P assisted in conducting experiments. P.Z. performed pathway enrichment analysis. Y.G. conceived the study and interpreted the data. M.J., A.X.K., and Y.G. wrote the manuscript. All authors contributed meaningful insights during discussions and reviewed and approved the final version of the manuscript.

## DATA AND MATERIALS AVAILABILITY

All data needed to evaluate the conclusions in the paper are present in the paper and/or the Supplementary Materials.

