## Supplementary Materials for "Proteome-wide reverse molecular docking reveals folate receptor as a mediator of PFAS-induced neurodevelopmental toxicity"

**This PDF file includes:**

Supplementary Figures 1 to 5

Supplementary Table 1 to 7 Legends

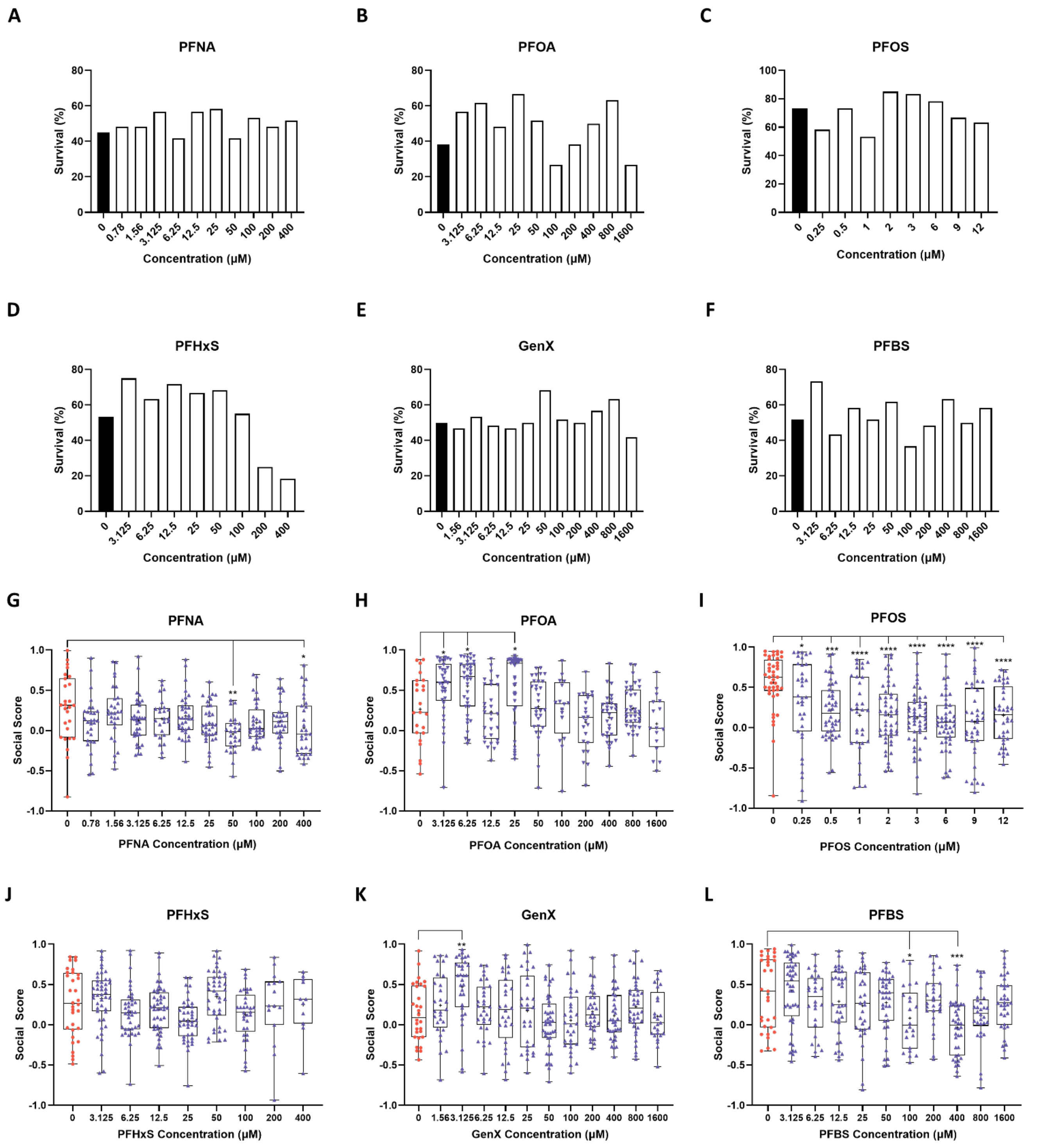


**Supplementary Figure 1.**

(A-F) The survival rates of all PFAS treatment conditions. Note that there is a ~50% natural attrition during the nursery growth period (5 dpf to 23 dpf), resulting in only ~50% survival in the vehicle controls (0 µM PFAS; black bars).

(G-L) Box plots of individual social scores (y-axis) for all PFAS treated fish, showing the distribution of social scores for each PFAS (G: PFNA, H: PFOA, I: PFOS, J: PFHxS, K: GenX, and L: PFBS) and concentration (x-axis). Significance was calculated by one-way ANOVA and Dunnett's multiple comparison test. Each date point shows the social score of an individual fish. $n \geq 16$ for all doses. *: $p < 0.05$, **: $p < 0.01$, ***: $p < 0.001$, ****: $p < 0.0001$.

A

| **PFNA-protein interactions with binding energy <=-10** | 27. TRAPPC3L | **PFOS-protein interactions with binding energy <= -10** | 31. ABHD6 |
|---|---|---|---|
| 1. CD1B | 28. SEC14L4 | 1. FOLR2 | 32. ST8SIA6 |
| 2. FOLR2 | 29. OR7C2 | 2. PLCB2 | 33. DTWD1 |
| 3. EBPL | 30. RFX3 | 3. SIRT1 | 34. SULT1A1 |
| 4. OR10P1 | 31. OR10AD1 | 4. WSCD1 | 35. CD1D |
| 5. OR10K1 | 32. AKR1C3 | 5. SLC7A6 | 36. FOLR3 |
| 6. CHST4 | 31. HS6ST1 | 6. CHST5 | 37. SULT1A2 |
| 7. OR6J1 | 32. LRP2 | 7. UGT2B28 | 38. COQ10A |
| 8. OR10AG1 | 33. SLC27A4 | 8. CD1C | 39. ST8SIA3 |
| 9. CD1A | 34. SLC6A14 | 9. KLHL10 | 40. TRAPPC3 |
| 10. OR8D2 | 35. PIGF | 10. NR1I3 | 41. AKR1C4 |
| 11. CHST5 | 36. SIGMAR1 | 11. NDUFA6 | 42. DHDDS |
| 12. PLCB2 | 37. SCARB2 | 12. ST8SIA2 | 43. CERT1 |
| 11. CHRM2 | 38. LRP2 | 13. SCARB2 | 44. PNP |
| 12. SCP2D1 | 39. MBOAT4 | 14. SCP2D1 | 45. NAA60 |
| 13. SLC7A6 | 40. ABHD6 | 15. AKR1C3 | 46. SULT2B1 |
| 14. AKR1B15 | 41. ELP1 | 16. LRP2 | 47. FOLH1 |
| 15. PTPN9 | 42. LCN9 | 17. LRP2 | 48. PIGF |
| 16. OR2L3 | 41. EBP | 18. CD1A | 49. DCAF5 |
| 17. OR5D18 | **PFOA-protein interactions with binding energy <=-10** | 19. AKR1B15 | 50. SEC14L4 |
| 18. PXMP2 | | 20. GBP3 | 51. EBP |
| 19. MPV17 | | 21. SULT1A4 | 52. CD1E |
| 20. LSS | 1. CD1B | 22. MPV17 | 53. ELP1 |
| 21. CERT1 | 2. OR10A7 | 23. KLHL36 | 54. ACSM5 |
| 22. RFX1 | 3. FOLR2 | 24. PTPN9 | 55. NAALAD2 |
| 21. GPR12 | 4. NIPA1 | 25. HHATL | 56. LRP2 |
| 22. RFX2 | 5. OR5D14 | 26. UROD | 57. QNG1 |
| 23. OR5A2 | 6. OR2L3 | 27. KDM4D | 58. MOSPD2 |
| 24. OR2G2 | 7. CHST6 | 28. LRP2 | 59. VWA7 |
| 25. NDUFA6 | 8. CHST5 | 29. TRMT10A | 60. ZDHHC5 |
| 26. FOLH1 | 9. AKR1B15 | 30. HSD17B13 | 61. SULT1B1 |

B

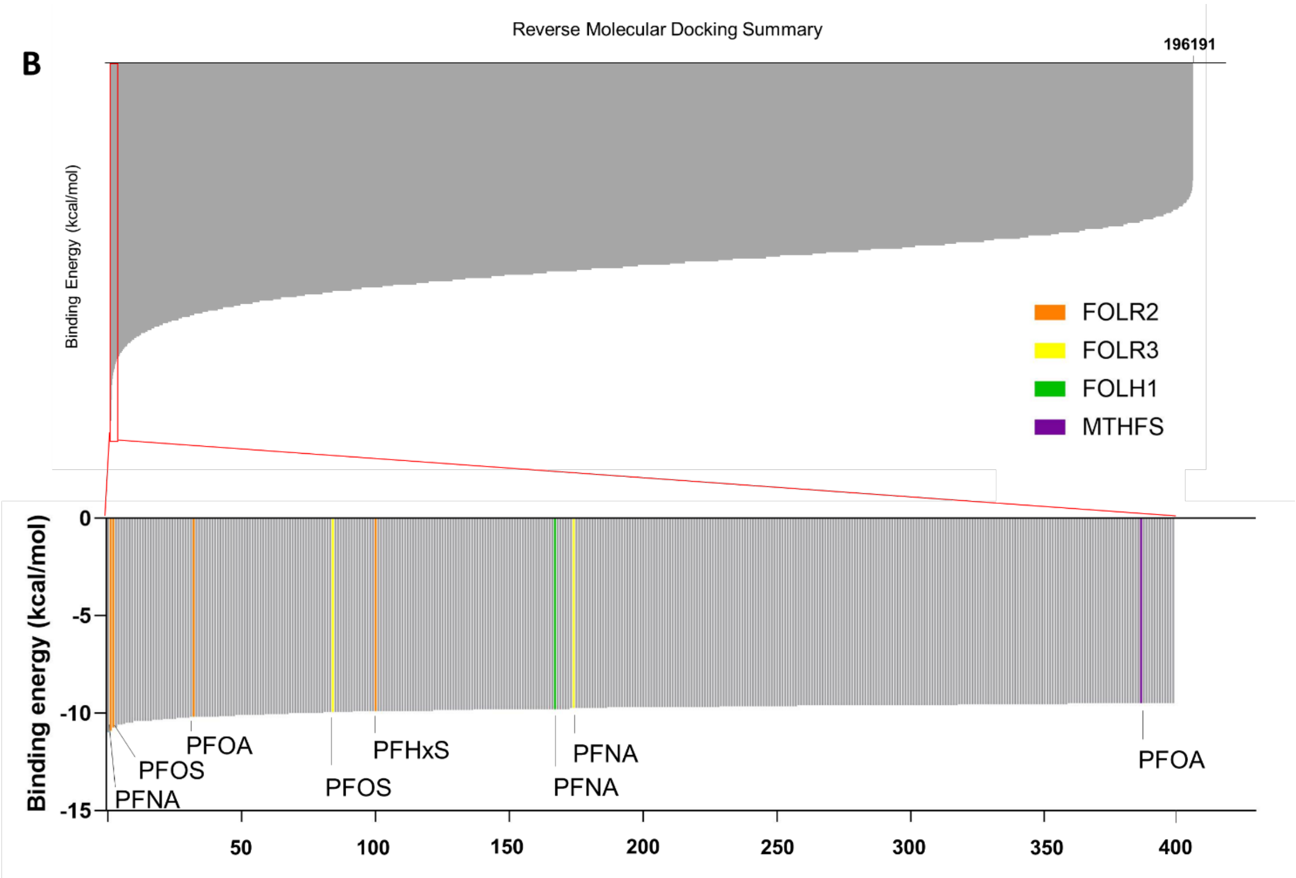

**Supplementary Figure 2.**

(A) The predicted protein targets of PFNA, PFOA, and PFOS. Hits were identified by selecting protein-PFAS interactions with predicted binding energy < -10 kcal/mol.

(B) Summary of the reverse molecular docking screen. All molecular docking scores of six EPA-regulated PFAS compounds against predicted binding pockets in 80% of human proteome are shown as a bar chart at the top. The top 400 highest affinity binding interactions (the lowest scores) are zoomed in and shown as a bar chart at the bottom. Interactions between PFAS (PFNA, PFOS, and PFOA) and FOLR2 (orange), FOLR3 (yellow), FOLH1 (green), and MTHFS (purple) are color-coded and highlighted. Ranking is based on average binding scores for each PFAS-domain interaction, calculated from binding energies between each PFAS and binding pockets predicted by structural-based prediction and sequence-based prediction for each protein domain.

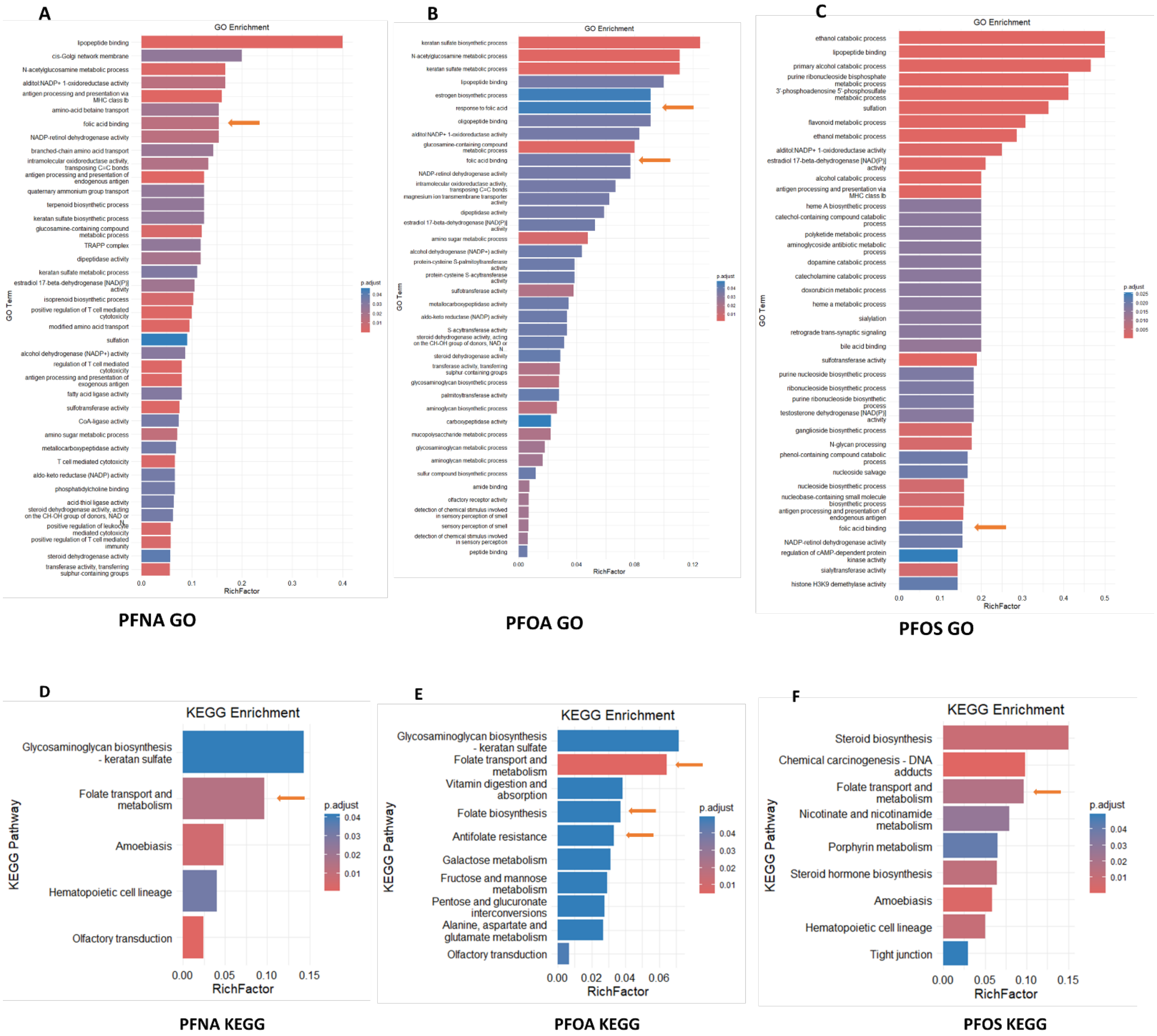


**Supplementary Figure 3.**

(A-C) Enrichment analysis for GO pathways consistently identified the folate pathway (orange arrows) for PFNA (A), PFOA (B), and PFOS (C).

(D-E) Enrichment analysis for KEGG pathways consistently identified the folate pathway (orange arrows) for PFNA (D), PFOA (E), and PFOS (F).

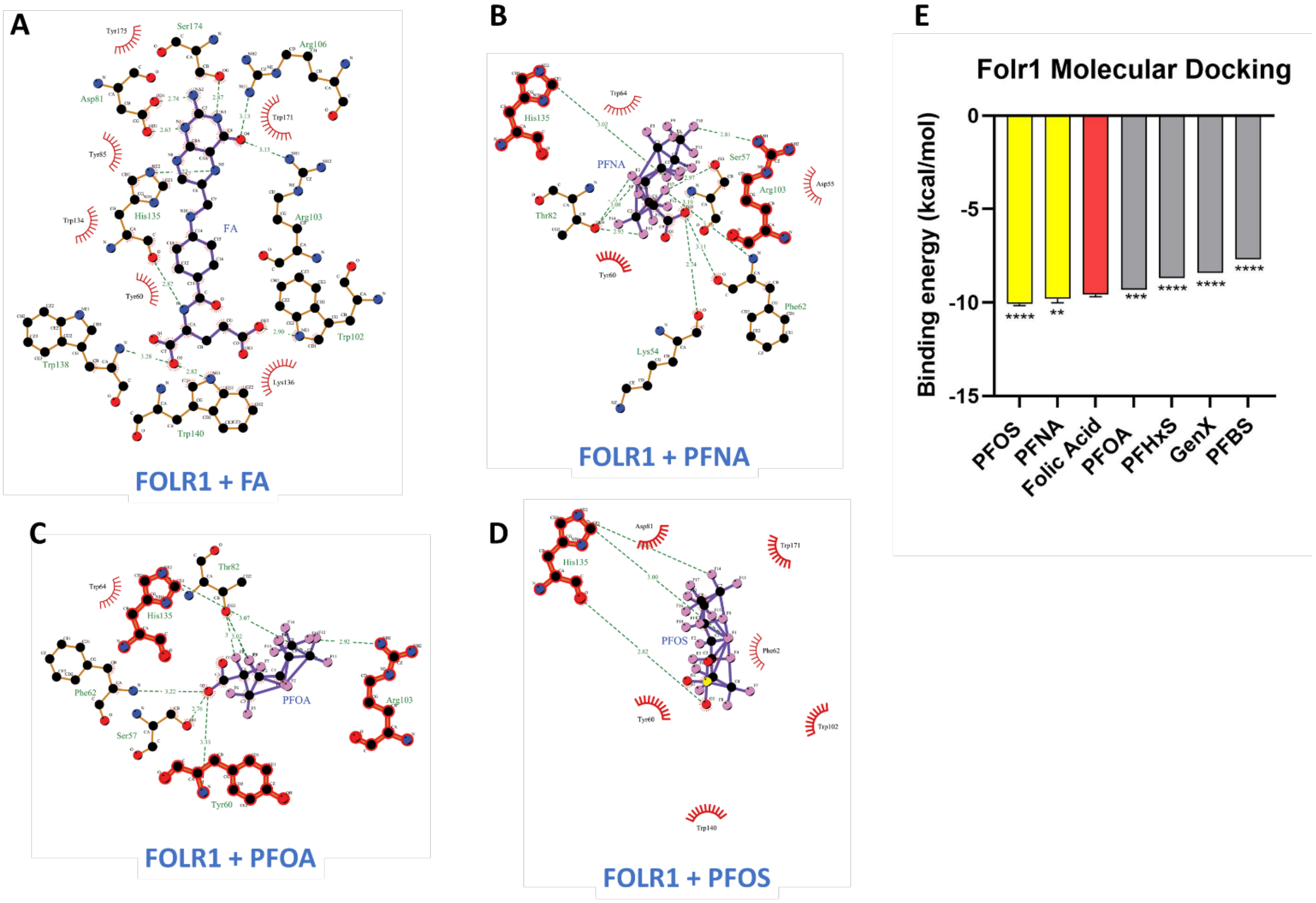


**Supplementary Figure 4.**

(A-D) *In silico* validation via molecular docking. Showing docking images for FOLR1 against folic acid (FA) (A), PFNA (B), PFOA (C), and PFOS (D). FOLR1 structure was acquired using the PDB identifier 4LRH.

(E) Comparing the binding energies for folic acid and the six EPA-regulated PFAS against FOLR1. Binding energies are shown as average and standard deviation of 5 independent docking attempts. Significance values are calculated to compare PFAS data with folic acid. Significance was calculated by one-way ANOVA and Dunnett's multiple comparison test. **: $p < 0.01$, ***: $p < 0.001$, ****: $p < 0.0001$.

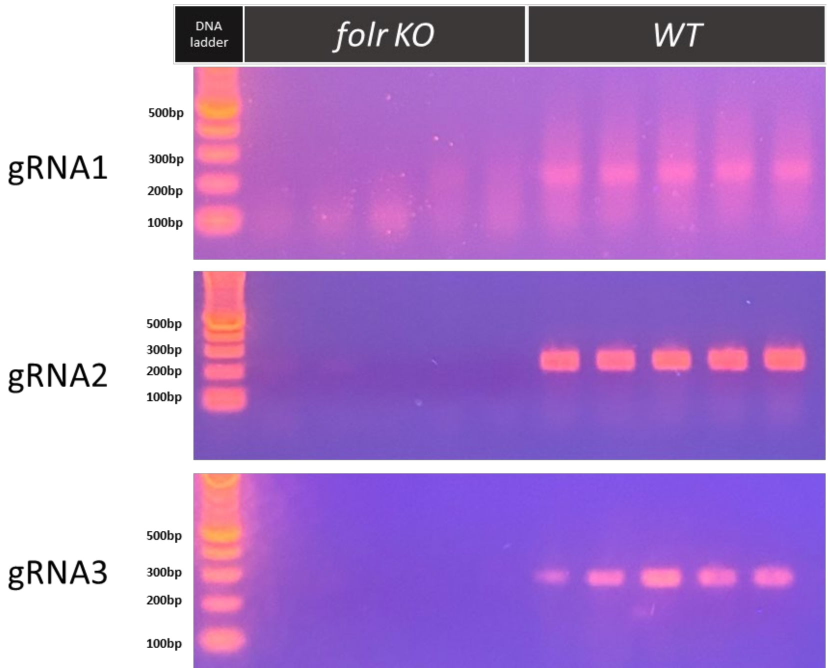


**Supplementary Figure 5.**

DNA gel analysis shows the disappearance and smearing of wild-type PCR amplification products for the genomic regions targeted by each *folr* gRNA following CRISPR/Cas9 knockout (KO). Five independently injected *folr* KO embryos and wild-type (WT) embryos were examined.

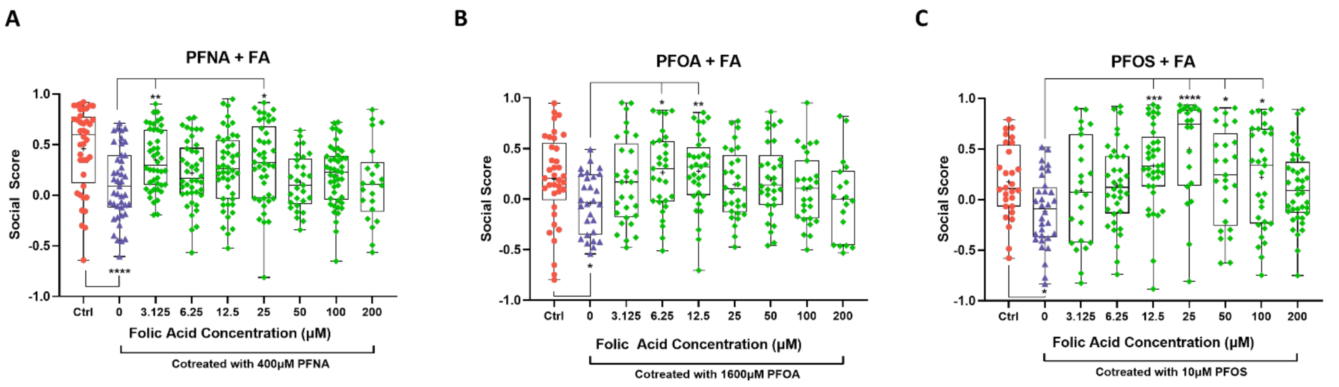


**Supplementary Figure 6.**

(A-C) Box plots of individual social scores (y-axis) for PFAS and folic acid (FA) co-treated fish, showing the distribution of social scores for each co-treatment (A: PFNA+FA, B: PFOA+FA, and C: PFOS+FA) and folic acid concentration (x-axis). Significance was calculated by one-way ANOVA and Dunnett's multiple comparison test. *: $p < 0.05$, **: $p < 0.01$, ***: $p < 0.001$, ****: $p <$ 0.0001.

### SUPPLEMENTARY TABLES

#### Supplementary Table 1.

Summary of the reverse molecular docking screen results for all 6 PFAS species.

#### Supplementary Tables 2-7.

Summary of the reverse molecular docking screen results for PFNA (Supplementary Table 2), PFOA (Supplementary Table 3), PFOS (Supplementary Table 4), PFHxS (Supplementary Table 5), GenX (Supplementary Table 6), and PFBS (Supplementary Table 7). Predicted binding affinities are ranked by binding energy from lowest (highest affinity) to highest (lowest affinity).
